# Soil health increases primary productivity across Europe

**DOI:** 10.1101/2023.10.29.564603

**Authors:** Ferran Romero, Maëva Labouyrie, Alberto Orgiazzi, Cristiano Ballabio, Panos Panagos, Arwyn Jones, Leho Tedersoo, Mohammad Bahram, Carlos A. Guerra, Nico Eisenhauer, Dongxue Tao, Manuel Delgado-Baquerizo, Pablo García-Palacios, Marcel G.A. van der Heijden

## Abstract

A healthy soil is at the core of sustainable management and policy, but its importance for plant productivity across environmental gradients and land-use types remains poorly understood. To address this gap, we conducted a pan-European field study including 588 sites from 27 countries to investigate the link between soil health and primary productivity across three major land-use types: woodlands, grasslands, and croplands. We found that mean soil health (a composite index based on soil properties, biodiversity, and plant disease control) in woodlands was 31.4% higher than in grasslands, and 76.1% higher than in croplands. Soil health was positively linked to cropland and grassland productivity at the continental scale. Woodland productivity was best explained by climate. Among microbial diversity indicators, we observed a positive association between the richness of Acidobacteria, Firmicutes, and Proteobacteria and primary productivity. Among microbial functional groups, we found that nitrogen-fixing bacteria and mycorrhizal fungi positively related to primary productivity in croplands and grasslands, while plant pathogens showed a negative relationship. Together, our results point to the importance of soil biodiversity and soil health for maintaining primary productivity across contrasting land-use types.

## Introduction

Plant productivity is fundamental for supporting key ecosystem services such as food provision and climate regulation (Barrios, 2007). A range of studies have shown that plant diversity in ecosystems promotes plant productivity and yield (Dietrich et al., 2023; Fraser et al., 2015; Isbell et al., 2011, 2015; Le Provost et al., 2023; Tilman et al., 1997). However, much less is known on how soil health including soil biodiversity relates to primary productivity, particularly across contrasting land-uses at large scales. Soil health refers to the capacity of soils to continuously support plant productivity and ecosystem functions (Lehmann et al., 2020; Qiao et al., 2022). Soil health assessments have traditionally focused on soil properties including microbial biomass, available nutrients, and organic carbon (Bünemann et al., 2018; Lehmann et al., 2020). However, the broader implications of integrating soil biodiversity into these assessments, especially when conducted across diverse land-uses and large spatial scales, remain underexplored yet potentially significant for ecosystem management.

Despite the growing interest in soil health and how it contributes to human well-being (Bender et al., 2016; Wall et al., 2015), current scientific evidence that demonstrates the importance of soil health at large scale is missing (Lehmann et al., 2020; Rinot et al., 2019). A recent study demonstrated that soil health is positively linked to plant yield in croplands in China (Li et al., 2021). Yet, the relative contribution of soil health to explain primary productivity across contrasting ecosystems (i.e., woodlands, grasslands, croplands) remains unknown. The mechanisms by which soil health drives productivity are many-fold: soil organic carbon, for example, positively contributes to primary productivity as it increases soil water retention and nutrient availability (Rawls et al., 2003). On the other hand, the presence and richness of plant pathogens in the soil can lead to diseases, reducing plant growth and yield (Maron et al., 2011). The link between soil health and plant productivity between different land-use types might differ due to variations in biological, physical, and chemical soil properties. Mature woodlands, for example, are stable ecosystems and thought to display high levels of soil health and are dominated by trees with deep root systems that can access nutrients from deeper soil layers and can store nutrients in their biomass for decades (Fernández-Martínez et al., 2014). Additionally, the slow decomposition of leaf litter in woodlands contributes to soil organic matter and nutrient availability over time, which may lead to a gradual increase in primary productivity as the forest matures (Krishna & Mohan, 2017). Consequently, as it is already high, soil health may be less important for productivity in woodlands and other factors (e.g., water availability, temperature) may have a bigger impact on woodland productivity. On the other hand, grasslands and croplands are subjected to more disturbance and are often characterized by dense and short root networks of grasses and forbs that grow and die seasonally, leading to a rapid turnover of nutrients and organic matter (Gill & Jackson, 2000; Wang et al., 2019).

Recently, it was observed that soil microbial diversity at the European scale is highest in croplands compared to other ecosystems, suggesting that diversity and plant productivity may be decoupled in these human-made ecosystems (Karimi et al., 2019; Köninger et al., 2023; Labouyrie et al., 2023; Szoboszlay et al., 2017). We also know that certain microbial groups are particularly beneficial for primary productivity, including nitrogen-fixing (N-fixing) bacteria and mycorrhizal fungi (Bauer et al., 2012; van der Heijden et al., 2015). However, the question of whether productivity is similarly driven by biological components of soil health (i.e., microbial diversity) across contrasting land-use types remains unexplored, as a systematic large-scale comparison among different land-uses is lacking and large-scale monitoring efforts are scarce (Guerra et al., 2021; Heintz-Buschart et al., 2020).

Here, we tested whether soil health is positively linked to primary productivity across Europe. In addition, we tested which soil variables best explained primary productivity. Our analysis is based on 588 sites from 27 countries, and we assessed three major land-use types in Europe: woodlands (n = 185), grasslands (n = 126), and croplands (n = 277). As a measure of soil health, we used a composite index built from seven soil properties including abiotic and biotic factors (i.e., soil organic carbon, phosphorus content, water infiltration potential, microbial biomass, richness of N-fixing bacteria, richness of mycorrhizal -arbuscular and ectomycorrhizal-fungi, and plant disease control - see methods). As a measure of primary productivity, we used the normalized difference vegetation index (NDVI) derived from remote sensing data obtained from each of the 588 sampling locations. Specifically, we tested whether the relationship between soil health and productivity differs among contrasting land-use types. We hypothesized that less disturbed ecosystems (i.e., woodlands) would show healthier soils compared to grasslands and croplands. We also hypothesized that primary productivity in grasslands and croplands would be more immediately impacted by soil health compared to woodlands.

## Methods

### Field survey and soil sampling

This study was carried out within the EU Statistical Office’s Land Use and Coverage Area frame Survey (LUCAS), the largest pan-European scheme for assessing soil characteristics in relation to land cover and use (Orgiazzi et al., 2018). As part of the 2018 LUCAS Soil survey, 881 fresh soil samples were collected across Europe and stored at −20°C before further processing (Orgiazzi et al., 2022). The sampling followed a composite strategy: the final sample collected at each location comprised five topsoil (0-20 cm) subsamples that are physically mixed to form a single composite sample. The first subsample was taken at the precise GIS (geographical information system) coordinates of the pre-established LUCAS point, and the remaining four subsamples were taken 2 m from the central one following the cardinal directions (North, East, South, and West). For this study, we focused on 277 cropland, 126 grassland, and 185 woodland sites across 27 countries (Figure 1). Croplands included both permanent (e.g., fruit trees, olive groves, vineyards) and non-permanent (e.g., cereals, legumes) sites. Grasslands included sites covered by communities of grasses, grass-like plants, or forbs. Woodlands included both broadleaved and coniferous forests. Sampling lasted from May to December 2018. The selection was based on availability of all critical variables (soil biodiversity, edaphic factors, and primary productivity).

**Figure 1.**
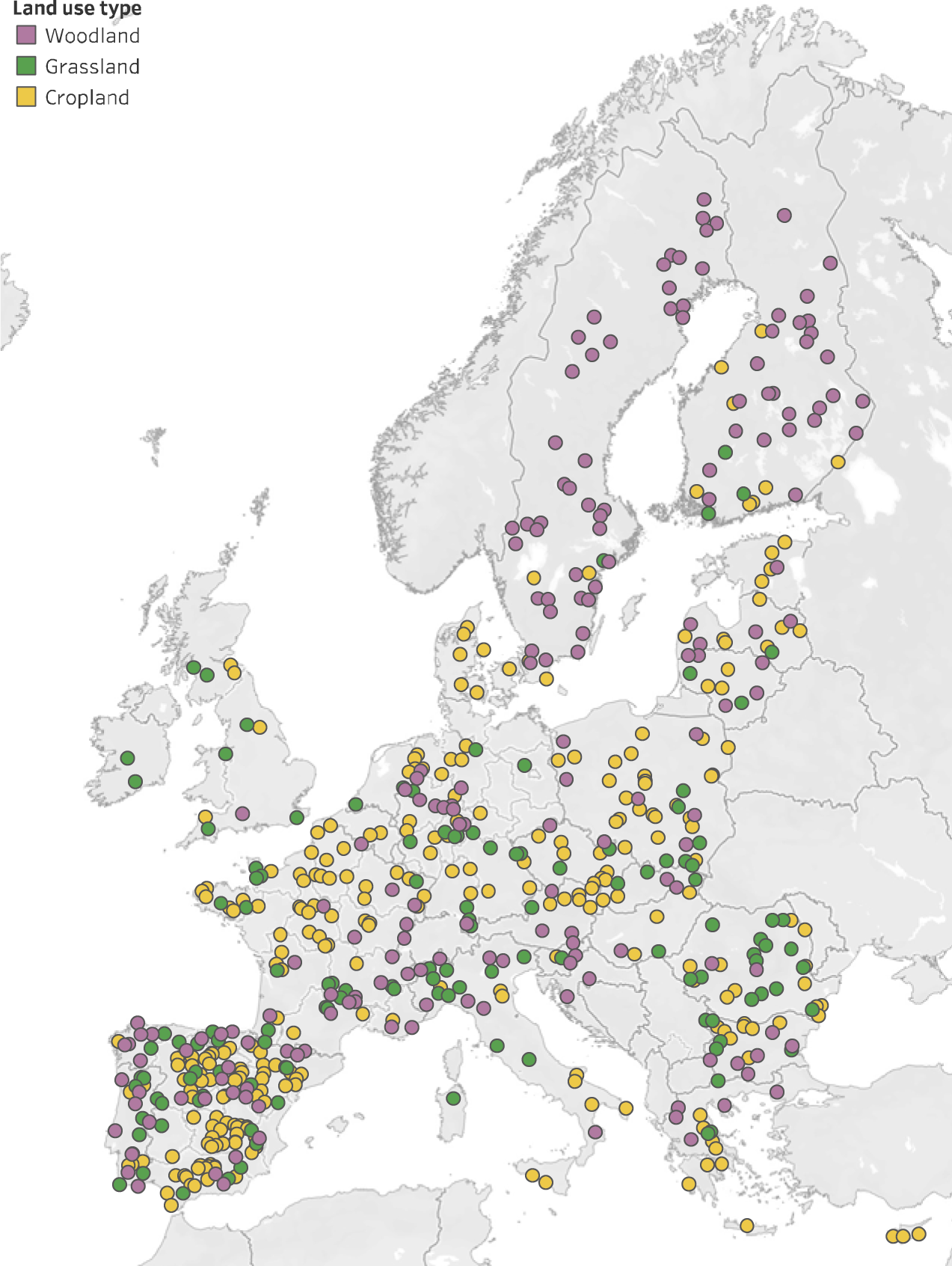
Study site locations. Geographic distribution across the European continent of the 588 locations used in this study. Three major land-use types were covered in the survey: cropland (n = 277, yellow), grassland (n = 126, green), and woodland (n = 185, purple).

### Soil microbial diversity

Soil DNA was extracted from each of the 588 composite soil samples using the Qiagen DNeasy PowerSoil HTP 96 Kit Q12955-4. Three 0.2 g aliquots per sample were extracted and pooled. Negative and positive controls were included to determine the presence of any external contamination. DNA extracts were quantified and checked for quality using a Qubit^TM^ 1X dsDNA HS Assay Kit and a Qubit^TM^ 3 Fluorometer (Invitrogen). Primer sets for barcode amplification were 515F (GTGYCAGCMGCCGCGGTAA) and 926R (GGCCGYCAATTYMTTTRAGTTT) for the bacterial 16S region (Caporaso et al., 2011; Parada et al., 2016), and ITS9mun (GTACACACCGCCCGTCG) and ITS4ngsUni (CGCCTSCSCTTANTDATATGC) for the fungal ITS region (Tedersoo & Anslan, 2019). Sequencing was performed by Illumina MiSeq platform with 2 x 300 paired-end mode for bacterial amplicons and PacBio Sequel II platform for fungal amplicons. Additional information including DNA amplification and library preparation is available in Labouyrie et al., 2023.

The Illumina and PacBio amplicon data (for bacteria and fungi, respectively) were demultiplexed using LotuS2 (Özkurt et al., 2022), and paired-end reads were assembled setting a minimum overlap of 10 bp using FLASH 1.2.10 (Magoč & Salzberg, 2011). Reads failing to merge were removed from the final dataset. For bacteria, operational taxonomic units (OTUs) were generated with the *usearch* option (version 10.0.024) available in UPARSE (Edgar, 2013). Merged reads were truncated up to the 16S primer sequences (i.e., 515F and 926R) and filtered for the presence of primer sequences (allowing a maximum of 2 mismatches). Merged reads were further quality-filtered with usearch by removing sequences with more than one expected error, and duplicated sequences were collapsed with *fqtrim* version 0.9.7 (Pertea, 2015). Obtained OTUs were annotated with the taxonomy data available from the Ribosomal Database Project (Cole et al., 2014). To avoid sequencing artifacts, only OTUs with at least 50 read counts and present in at least 10 samples were kept. For fungi, 98%-OTUs were obtained as described in Tedersoo et al., 2021. Fungal taxonomic annotation was performed using BLAST+ version 2.11.0 (Camacho et al., 2009) against the UNITE 9.1 database (Nilsson et al., 2019). Bacterial and fungal datasets were further rarefied to comparable sequencing depths (bacteria: 40,109 reads; fungi: 502 reads). Rarefied datasets were used to obtain values of total richness (i.e., OTU number), community composition (extracted values from the first dimension of NMDS based on Bray-Curtis dissimilarities), and the richness of major microbial groups (i.e., phyla). Rarefied datasets were also used to obtain the richness of major functional groups as retrieved from the FAPROTAX (bacteria) and FungalTraits (fungi) databases (Louca et al., 2016; Põlme et al., 2020) (Table S1). Briefly, we retrieved the number of OTUs classified as bacterial chemoheterotrophs, nitrogen-fixing bacteria, fungal saprotrophs, fungal plant pathogens, and mycorrhizal fungi, as described in Liu et al., 2022. Further details on bacterial/fungal sequence processing, filtering, and taxonomic/functional annotation are available in Labouyrie et al., 2023.

### Edaphic and climatic factors

The following measurements were taken for each sampling site to ensure a comprehensive understanding of climatic and edaphic factors (Ballabio et al., 2019): Total nitrogen and phosphorus content were assessed to determine nitrogen and phosphorus availability in the soil. Soil organic carbon was quantified to estimate the amount of carbon stored in the soil organic matter. Soil pH and microbial biomass were used as measures of acidity and activity of the living component of the soil, respectively. Soil bulk density was used as a surrogate of water infiltration potential. Soil texture was characterized by measuring silt, sand, and clay relative content. Additionally, to capture the broader climatic context of each site, mean annual precipitation and mean annual temperature (i.e., averaged monthly values over a long period of time; 1970-2000) were retrieved from the WorldClim database (www.worldclim.org). These parameters collectively provided a detailed profile of the environmental and soil conditions at each sampling site (Table S1). Specific details on the methods employed for these measurements are available in Labouyrie et al., 2023, Orgiazzi et al., 2018 and Smith et al., 2021.

### Primary productivity

The Net Primary Productivity (NPP) was derived from images taken by the NASA Terra and Aqua satellites using their MODerate resolution Imaging Spectroradiometers (MODIS). The NPP is calculated as the sum of the MODIS eight-days Net Photosynthesis (PSN) products. PSN is in turn estimated from the Fraction of Absorbed Photosynthetically Active Radiation (FAPAR,), which is the fraction of the incoming solar radiation (in the photosynthetically active spectrum) absorbed by plants. FAPAR is closely related to NDVI (normalized difference vegetation index), that has been used as a proxy for plant biomass in several studies relating soil biodiversity and primary productivity (Delgado-Baquerizo et al., 2016; Fan et al., 2023; Liu et al., 2022). However, the derivation of PSN from FAPAR, provides a more reliable estimate of the NPP as it accounts for the ground conditions in which the photosynthesis occurs and for the plant maintenance respiration. Thus, this computation requires additional information such as the maximum radiation conversion efficiency, the ground temperature, and vapour pressure. The calculation is performed following the MOD17 PSN/NPP algorithm (Running et al., 1999). For this study the NPP was derived from MODIS products for year 2018 (to match with the LUCAS survey) at a resolution of 500 m (as described in Ballabio et al., 2016, 2019). The NPP values were derived for the months before, during, and after sampling at each LUCAS location and subsequently averaged.

### Soil health

In this study, we used seven explanatory variables (selection based on Lehmann et al., 2020), namely soil organic carbon, phosphorus content, water infiltration potential, microbial biomass, richness of N-fixing bacteria, richness of mycorrhizal fungi, and plant disease control) to build a soil health index (Table S2). These variables were selected as they collectively represent a broad range of soil characteristics that are critical for understanding soil health-productivity relationships. Soil organic carbon is crucial for nutrient cycling and soil structure; phosphorus content is a key nutrient for plant growth; water infiltration potential influences water availability and root development; microbial biomass indicates the biological activity essential for nutrient transformation; richness of N-fixing bacteria and mycorrhizal fungi reflects soil capacity for nitrogen fixation and nutrient uptake, respectively; and plant disease control highlights the role of soil in supporting plant health. The soil health index was calculated using z-score transformation: each variable was standardized to have a mean of 0 and a standard deviation of 1. The soil health index for each site was then computed by summing its respective z-scores (i.e., equally weighting). This approach allowed us to integrate multiple soil health indicators into a single composite index, facilitating a comprehensive assessment.

### Statistics

Statistical analyses were performed using R software, version 4.2.1. Only soil samples for which all explanatory variables (i.e., soil biodiversity and climatic, edaphic, and spatial factors) were available were used for this study, so none of the values in any explanatory variables was imputed or calculated from the median value of the respective dataset. A total of 588 sites were used for this study, including 277 croplands, 126 grasslands, and 185 woodlands (Figure 1). To assess the most important predictors of primary productivity within the three land-use systems, we used Random Forest (RF) (Cutler et al., 2007) followed by structural equation modeling (Eisenhauer et al., 2015).

First, we selected a wide range of explanatory variables including biodiversity indicators (i.e., OTU richness, diversity, community composition), together with climatic and edaphic factors. The total number of explanatory variables was 34 (Table S1). We then investigated the correlation among all variables to ensure that there was no strong collinearity and reduced the variables that were strongly correlated (i.e., Spearman rank correlation coefficient higher than +0.70, or lower than −0.70). Correlations were run using the *corr* function within the *corrplot* package (Wei & Simko, 2017). This approach reduced the 34 explanatory variables to 24 (croplands), 23 (grasslands), and 19 (woodlands) (Table S1). The variables removed included alpha and beta diversity indices (strong correlation with specific microbial groups, e.g., Proteobacteria), total nitrogen (strong correlation with soil organic carbon), richness of chemoheterotrophic bacteria (strong correlation with Actinobacteria and total bacteria), and silt and sand content (strong correlation with clay content).

To further clarify the relative contribution of climate, edaphic factors, and soil biodiversity to explain patterns in primary productivity, we conducted RF analyses. The RF analyses were performed independently for the three land-use types using 10,000 permutations. RF analyses were performed using the *rfPermute* package (Archer, 2016). Predictor importance was estimated by the percent increase in Mean Squared Error (MSE) when each predictor was permuted. This reflects the contribution of each predictor to model accuracy; a higher increase in MSE indicates greater importance for accurate predictions. Finally, we employed Structural Equation Modelling (SEM) to evaluate the direct and indirect relationships between climate, edaphic factors, soil biodiversity, and primary productivity. We adapted the procedure described in Delgado-Baquerizo et al., 2016 using the R package *piecewiseR* for SEM (Lefcheck, 2016).

## Results and discussion

Here we assessed the link between soil health and primary productivity, examining 588 sites across three land-use types (i.e., croplands, grasslands, and woodlands) in Europe (Figure 1). We found the highest soil health values in woodlands compared to grasslands and croplands. According to our initial hypothesis, soil health in woodlands was ∼31% higher than in grasslands, and ∼76% higher than in croplands (Figure 2). Soil health was positively linked to primary productivity when all sites were taken together (Figure 2). In line with our expectations, soil health was positively linked to primary productivity in grasslands and croplands, but not in woodlands (Figure 2). Our soil health index encapsulated a range of abiotic and biotic soil factors, thus providing a comprehensive assessment of soil quality. This approach is advantageous as it represents a holistic measure that might serve as a decision-making tool for farmers and land managers. In our study, utilizing a comprehensive soil health index resulted in a stronger correlation with primary productivity than any single variable alone. Specifically, the correlation between soil health and productivity was 3.17 times higher than the average of individual variables for croplands, 2.64 times higher for grasslands, and 1.38 times higher for woodlands (Table S3).

**Figure 2.**
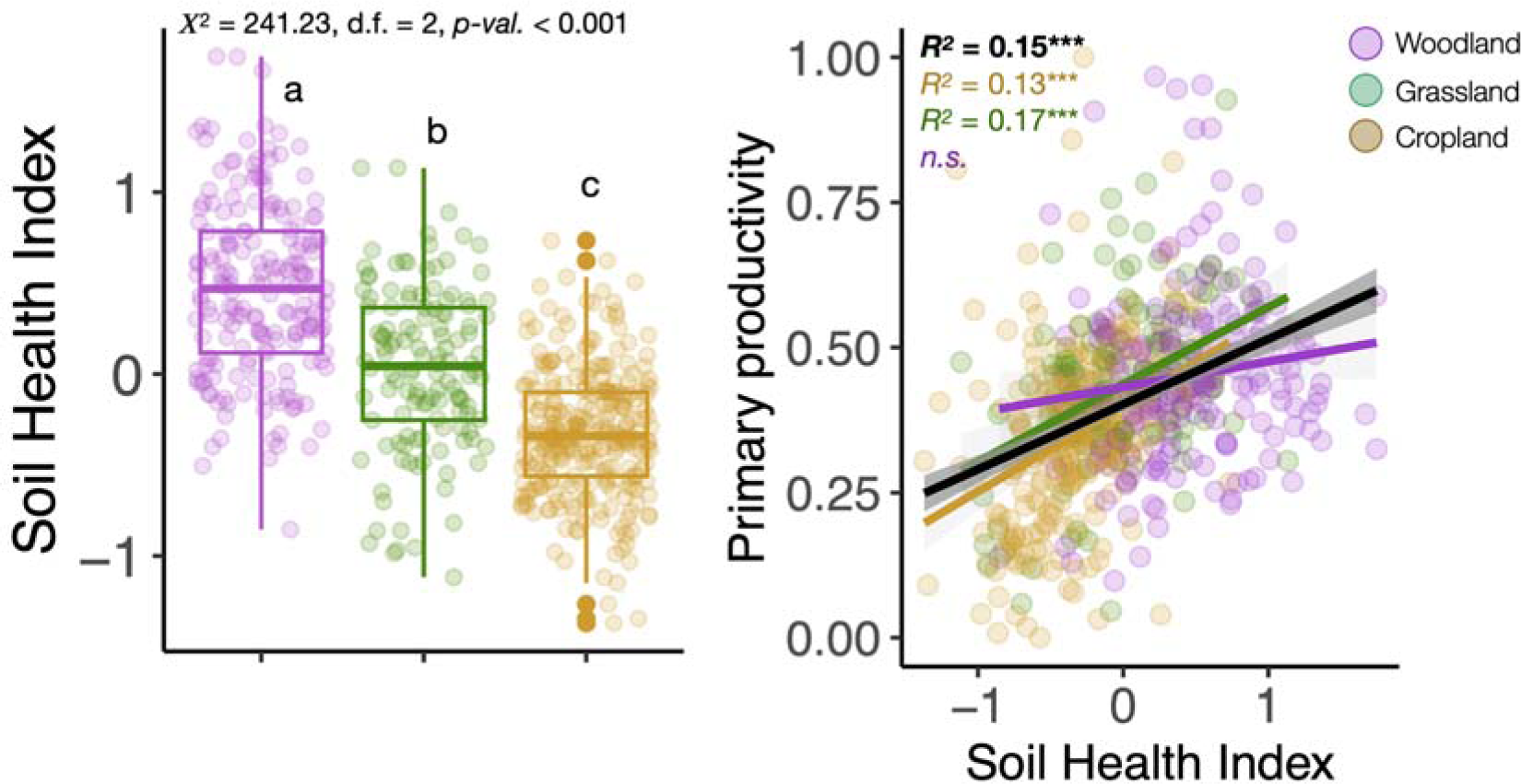
Relationship between soil health and primary productivity. Soil health index across land-use types (woodlands, grasslands, and croplands), and relationship between the soil health index and primary productivity. Explained variation is expressed as R^2^ (0-1), and significance is indicated by asterisks (***; p-value < 0.001, **; p-value < 0.010, *; p-value < 0.050). Non-significant (p-value > 0.05) regressions are denoted with “n.s.”.

Next, we assessed which individual variables best explained primary productivity. Overall, our results demonstrate that a combination of climatic and edaphic factors, together with soil biodiversity, explains patterns in primary productivity across contrasting land-use types at the European scale: our structural equation models explained 40% of the variability in primary productivity in grasslands, 35% in woodlands, and 31% in croplands (Figures 3 and 4). Our results demonstrate that primary productivity in cropland and grassland is best explained by various factors of soil health, including edaphic properties (e.g., organic carbon) and soil microbial diversity. In woodlands, our results show that temperature is a major regulator of primary productivity, although specific components of soil health (e.g., phosphorus content, organic matter, and the richness of Actinobacteria and Proteobacteria) also do play a role. Moreover, we observed that various components of microbial diversity act as relevant predictors of primary productivity, including the richness of Acidobacteria, Firmicutes, Proteobacteria, and mycorrhizal fungi (Figure 3, Figure 4).

**Figure 3.**
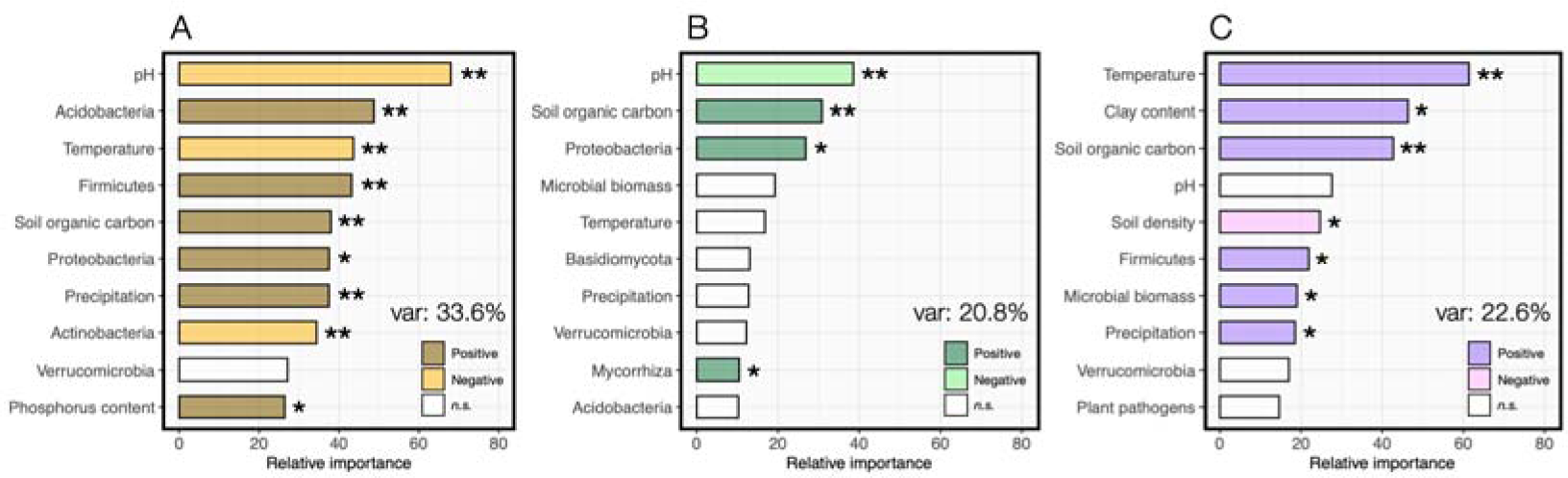
Relative importance of the main predictors of primary productivity. Relative importance of main predictors (top-10) for primary productivity based on random forest analysis in croplands (A, n = 277), grasslands (B, n = 126) and woodlands (C, n = 185). Dark colors indicate positive predictor effects on primary productivity, and light colors indicate negative effects. Variance explained by models is indicated as percentage (%). Asterisks indicate a significant effect based on the averaged model coefficients (***; p-value < 0.001, **; p-value < 0.010, *; p-value < 0.050).

**Figure 4.**
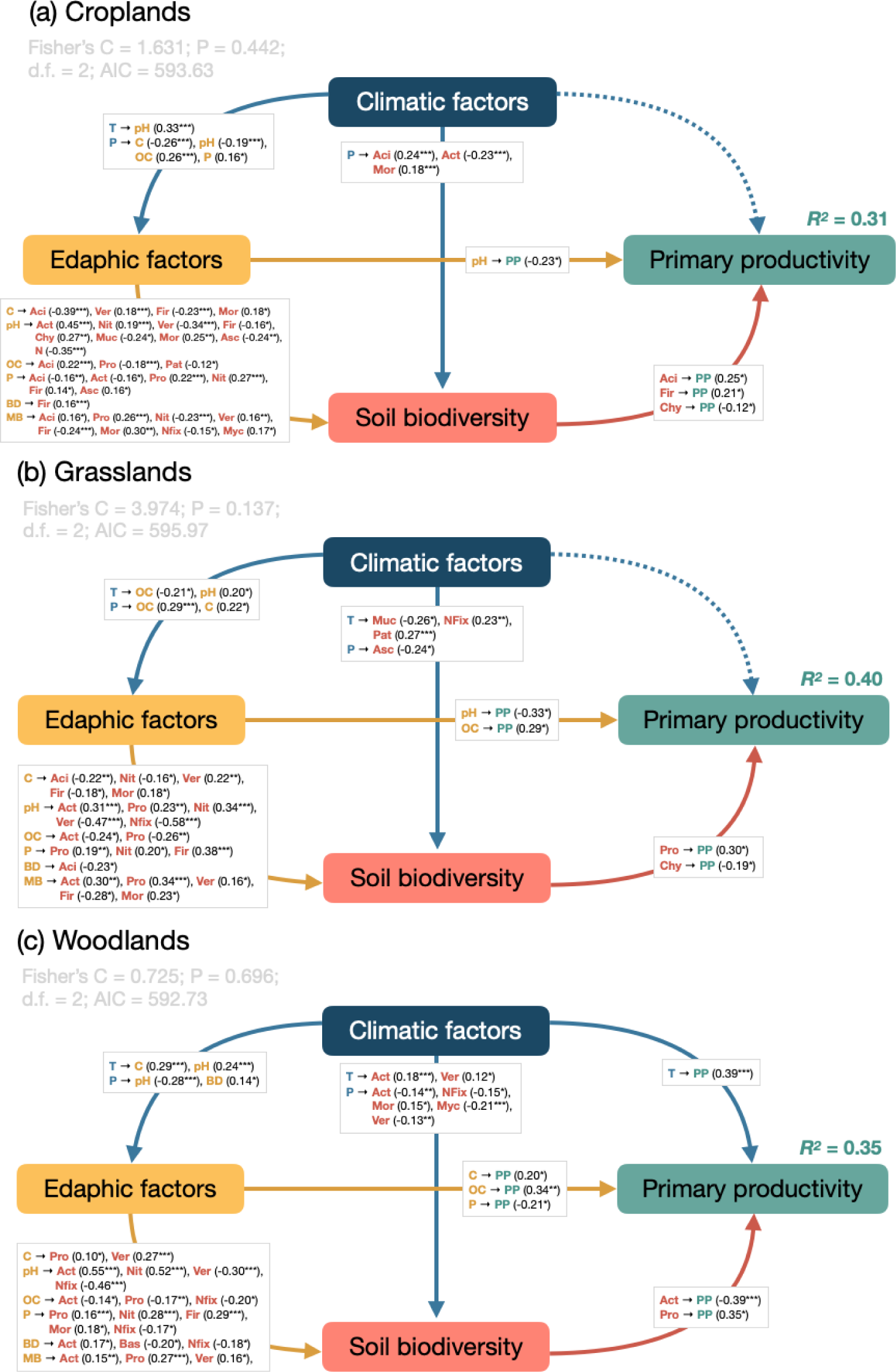
Structural equation modeling (SEM) describing the relationships between climate, edaphic factors, soil biodiversity, and primary productivity. Direct and indirect effects of climate, edaphic factors, and soil biodiversity on primary productivity across different land-use types. Variables included in each category (i.e., climatic factors, edaphic factors, and soil biodiversity) are indicated on the right panels. The proportion of explained variance (*R*^2^) appears alongside primary productivity in each model. Goodness-of-fit statistics are also included. Dashed lines indicate that none of the variables included was significant (p-value > 0.05). d.f.: degrees of freedom, AIC: Akaike information criterion. Variables included within “Edaphic factors” are pH, clay content (C), soil organic carbon (SOC), phosphorus content (P), soil bulk density (BD), and microbial biomass (MB). Variables included within “Climatic factors” are monthly air temperature (T) and monthly precipitation (P). Variables included within “Soil biodiversity” are Acidobacteria (Aci), Actinobacteria (Act), Mortierellomycota (Mor), Verrucomicrobia (Ver), Firmicutes (Fir), Nitrospirae (Nit), Chytridiomycota (Chy), Mucoromycota (Muc), Ascomycota (Asc), Basidiomycota (Bas), Proteobateria (Pro), N-fixing bacteria (Nfix), Mycorrhiza (Myc), and plant pathogens (Pat). “PP” stands for primary productivity.

Overall, our results indicate a positive relationship between soil health and cropland productivity (Figure 2). Random Forest analyses further indicated that pH was the main predictor of primary productivity in croplands. This confirms previous evidence on the role of pH in driving crop productivity (Couto, 2018). Accordingly, we observed the lowest cropland productivity in highly alkaline (pH ≈ 8) soils (Figure S1). Other edaphic factors showed positive effects on cropland productivity, namely soil organic carbon and phosphorus content (Figure 3, Figure S1). Among climatic factors, we observed a negative effect of increasing temperature on cropland productivity, while the effect of increasing precipitation was positive (Figure 3, Figure S1). Our results further indicate that microbial community composition was the best microbial predictor of cropland productivity (Figure S2). In line with this, random forest analyses also indicated that specific microbial groups predicted cropland productivity: that was the case for Acidobacteria, Firmicutes, and Proteobacteria. The richness of these bacterial groups was positively correlated with productivity, while a negative effect was observed for Actinobacteria (Figure 3, Figure S3). Structural equation models (SEMs) further demonstrated that edaphic factors (e.g., soil texture, soil organic carbon, pH, phosphorus content) affected cropland productivity also indirectly by leading to changes on the diversity of soil microorganisms, particularly the bacterial taxa Acidobacteria and Firmicutes, and the fungal phylum Chytridiomycota (Figure 4). Structural equation models confirmed a positive direct effect of increased richness of Acidobacteria and Firmicutes on cropland productivity, while the opposite was observed for Chytridiomycota (i.e., increased OTU richness led to lower productivity). Acidobacteria are abundant soil-borne microorganisms known to be involved in nutrient mineralization and organic matter degradation (Banerjee et al., 2016; Kielak et al., 2016; Sikorski et al., 2022). Our results also align with a recent study where Acidobacteria positively correlated with cropland productivity in a four-decade fertilization experiment (Fan et al., 2021). Similarly, members of the phylum Firmicutes, known for their widespread presence in soils and for promoting plant growth locally, appear to also enhance cropland productivity on a continental scale (Fierer, 2017; Hashmi et al., 2020). Finally, SEMs indicated that the richness of Chytridiomycota was negatively related to cropland productivity (Figure 4). While this fungal group is best known for parasitizing vertebrates, some species are responsible for plant disease in Europe (van de Vossenberg et al., 2022). In line with this, we observed a negative correlation between the richness of fungal plant pathogens and cropland productivity (Figure S3).

The grasslands included in this study were, on average, slightly more acidic than croplands and showed higher microbial biomass and higher organic carbon content, but lower phosphorus content compared (see full details in Table S4). Bacterial richness was lower in grasslands compared to croplands, but fungal richness was slightly higher (Table S4). As for croplands, our results indicate a positive relationship between soil health and grassland productivity (Figure 2). Random forest analyses retained four factors as predictors of primary productivity in grasslands, namely pH, soil organic carbon, and the richness of Proteobacteria and mycorrhizal fungi (Figure 3). Similar to croplands, pH showed a hump-shaped to negative response, with productivity peaking at pH ≈ 6. The relationship between grassland productivity and soil organic carbon, Proteobacteria, and mycorrhizal fungi was positive. Grasslands store one third of the terrestrial carbon stock, and most of it is stored belowground as soil organic carbon (Conant, 2012). Recent studies show that plant diversity increases soil organic carbon storage in grasslands by elevating carbon inputs to belowground biomass and promoting microbial necromass contribution to soil organic carbon storage (Lange et al., 2015; Xu et al., 2020). Our results add to this and show that soil organic carbon is positively linked to plant productivity for grasslands at the continental scale, indicating that soil organic carbon favors carbon storage, but also has positive effects on productivity. Although we did not differentiate between necromass and living biomass in our study, we also found positive correlations between bacterial richness/ microbial biomass and grassland productivity (Figure S1, Figure S2). Structural equation models further supported the idea that grassland productivity was determined by pH and soil organic carbon (Figure 4). Moreover, SEMs indicated a direct positive effect of Proteobacteria richness on grassland productivity, and a negative direct effect of Chytridiomycota.

Finally, woodlands showed the lowest pH and phosphorus content compared to croplands and grasslands, but the highest organic carbon content. Woodlands also showed the lowest values of bacterial and fungal richness (see full details in Table S4). Random forest modelling of the 185 woodland sites included in this study revealed that the best predictors of woodland productivity were monthly temperature, clay content, soil organic carbon, bulk density, richness of Firmicutes, microbial biomass, and monthly precipitation (Figure 3). Only bulk density showed a negative effect on woodland productivity, all other predictors were positively associated to it. SEMs revealed that woodland productivity was shaped by a complex interplay between climatic factors (i.e., temperature), edaphic factors (i.e., clay content, soil organic carbon, and phosphorus content) and soil biodiversity (richness of Actinobacteria and Proteobacteria). Specifically, SEMs depicted a strong positive effect of temperature on woodland productivity, although factors associated to soil diversity (e.g., richness of Actinobacteria and Proteobacteria) also shaped woodland productivity (Figure 4). Recent studies suggest that fungal communities are major drivers of ecosystem functioning in woodlands (Anthony et al., 2022; Osburn et al., 2023). Our SEM adds to this that fungal communities (particularly plant-beneficial members within Mortierellomycota, Basidiomycota, and mycorrhizal fungi) in woodlands are shaped by a combination of climatic factors (i.e., precipitation) and edaphic properties. Importantly, we also show that future studies addressing the impacts of the soil microbiome on woodland productivity need to include bacteria. This is because major bacterial divisions (i.e., Actinobacteria and Proteobacteria) were closely related to primary productivity in the woodlands included in our study.

In our study, primary productivity was significantly lower in croplands, compared to grasslands and woodlands, following previous research (Krause et al., 2022) (Figure S4). In line with our hypothesis, primary productivity relied more strongly on soil health in grasslands and croplands than in woodlands. Woodland productivity was largely governed by temperature, in line with previous findings (Collalti et al., 2020; Morin et al., 2018). Moreover, we observed the highest soil health values in woodlands. We argue that above a tipping point of soil health, other factors (i.e., climate), might become more important when it comes to explaining primary productivity. Finally, agricultural crops often have a short lifespan, rely on a fast delivery of nutrients and might be more dependent on soil health compared to less dynamic and more stable ecosystems like woodlands. In line with this, a recent study by Walder et al., 2023 assessed soil health and crop productivity in 60 fields managed either conventionally, under no-till, or organically. That study found that the management type with the lowest soil health, conventionally managed croplands, relied most on soil health when it comes to explaining plant productivity. Similarly, a recent greenhouse experiment showed that soil health is positively linked to nitrogen use efficiency (Toda et al., 2023). This indicates that soil health is especially important as driver of plant productivity and other ecosystem processes in disturbed agricultural soils. Among the different soil properties here assessed, we found strong correlations between total nitrogen and phosphorus and plant productivity across land-use types. We highlight that future studies should focus on their inorganic forms, as these are more readily available to plants in soil. This aspect was not addressed in our study due to the high number of samples involved.

Our study being based on soil health indices, we acknowledge that the diversity of soil systems makes it challenging to capture soil complexity into a single metric. Also, the assessment of soil health depends on the number of soil variables available, which is especially difficult for large scale assessments such as in this study. Here, we observed significant relationships between soil health and primary productivity, pointing to the importance of soil health as driver of plant productivity. However, these data are based on correlations, and further experimental studies need to complement and validate these findings empirically testing causality at specific locations (e.g., testing whether an increase in soil health indeed promotes productivity). In line with this, previous studies have successfully used both laboratory-based approaches and manipulative field studies to support findings from observational approaches (Delgado-Baquerizo et al., 2017; Sünnemann et al., 2023). Finally, we highlight the need to complement our NDVI-based vegetation analysis with actual plant biomass measurements, although we recognize the considerable technical challenges associated with conducting such detailed biomass studies at large scales.

In summary, our results provide compelling evidence that primary productivity is linked to soil health at the continental scale, particularly for grasslands and croplands. Among the soil health indicators assessed here, we highlight the positive effects of soil organic carbon and the richness of Acidobacteria, Firmicutes, and Proteobacteria on primary productivity. In contrast the richness of Actinobacteria and major plant pathogens including Chytridiomycota was negatively related primary productivity. Our study points to the importance of soil health as a driver of primary productivity and it highlights the need to include microorganisms in soil monitoring schemes to gain a comprehensive understanding of ecosystem functioning in terrestrial ecosystems.

## Acknowledgements

MvdH and FR acknowledge the Swiss National Science Foundation for funding through grant number 310030-188799. This project has received funding from the European Unionś Horizon 2020 research and innovation programme under grant agreement No. 862695 EJP SOIL-MINOTAUR. NE acknowledges funding by the Deutsche Forschungsgemeinschaft DFG (German Centre for Integrative Biodiversity Research, FZT118; and Gottfried Wilhelm Leibniz Prize, Ei 862/29-1; Ei 862/31-1). M.D-B. acknowledges support from TED2021-130908B-C41/AEI/10.13039/501100011033/Unión Europea NextGenerationEU/PRTR and from the Spanish Ministry of Science and Innovation for the I⍰+⍰D⍰+⍰i project PID2020-115813RA-I00 funded by MCIN/AEI/10.13039/501100011033. The LUCAS Survey is coordinated by Unit E4 of the Statistical Office of the European Union (EUROSTAT). The LUCAS Soil sample collection is supported by the Directorate-General Environment (DG-ENV), Directorate-General Agriculture and Rural Development (DG-AGRI) and Directorate-General Climate Action (DG-CLIMA) of the European Commission.

