## Supplementary Information for "Soil health increases primary productivity across Europe"

Ferran Romero*^1^, Maëva Labouyrie^1,2,3^, Alberto Orgiazzi^3,4^, Cristiano Ballabio^3^, Panos Panagos^3^, Arwyn Jones^3^, Leho Tedersoo^5^, Mohammad Bahram^6,7^, Carlos A. Guerra^8,9^, Nico Eisenhauer^8,9^, Dongxue Tao^10,11^, Manuel Delgado-Baquerizo^10^, Pablo García-Palacios^2,12^, Marcel G.A. van der Heijden*^1,2^

^1^Plant-Soil Interactions group, Agroscope, Reckenholzstrasse 191, 8046 Zurich, Switzerland

^2^Department of Plant and Microbial Biology, University of Zurich, Zollikerstrasse 107, 8008 Zurich, Switzerland

^3^European Commission, Joint Research Centre Ispra (JRC Ispra), Via Enrico Fermi 2749, 21027 Ispra, Italy

^4^European Dynamics, Brussels, Belgium

^5^Mycology and Microbiology Center, University of Tartu, Liivi 2, 50409 Tartu, Estonia

^6^Department of Botany, Institute of Ecology and Earth Sciences, University of Tartu, Liivi 2, 50409 Tartu, Estonia

^7^Department of Ecology, Swedish University of Agricultural Sciences, Uppsala, Sweden

^8^German Center for Integrative Biodiversity Research (iDiv) Halle-Jena-Leipzig, Leipzig, Germany

^9^Institute of Biology, Leipzig University, Leipzig, Germany

^10^Laboratorio de Biodiversidad y Funcionamiento Ecosistémico. Instituto de Recursos Naturales y Agrobiología de Sevilla (IRNAS), CSIC, Av. Reina Mercedes 10, E-41012, Sevilla, Spain.

^11^Institute of Grassland Science, Key Laboratory of Vegetation Ecology of the Ministry of Education, Jilin Songnen Grassland Ecosystem National Observation and Research Station, Northeast Normal University, Changchun 130024, China

^12^Instituto de Ciencias Agrarias, Consejo Superior de Investigaciones Científicas, Madrid, Spain

*Authors for correspondence: Ferran Romero and Marcel van der Heijden

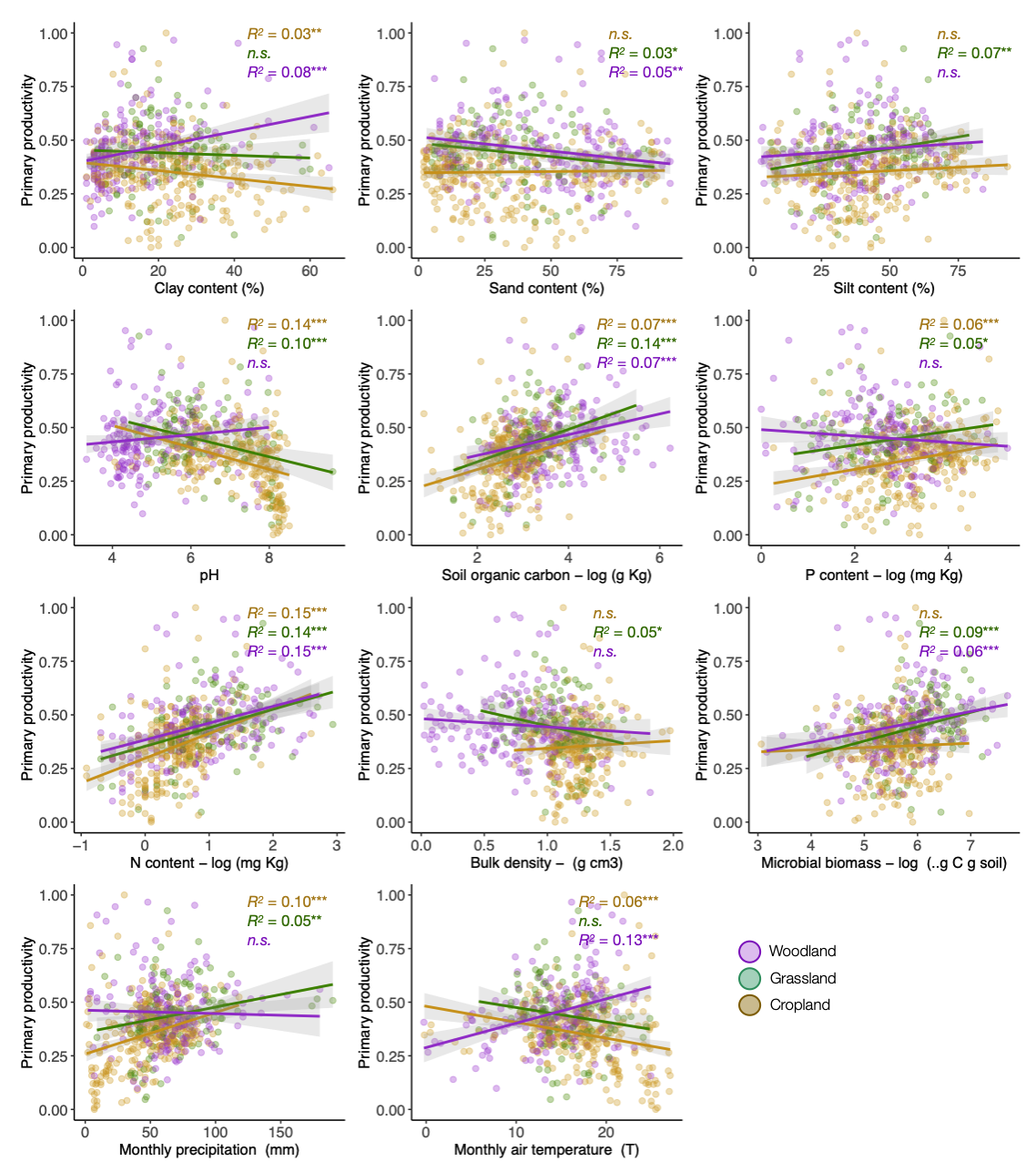

**Figure S1.** **Relationships between abiotic factors and primary productivity.** Linear regressions were fitted to explore the relationship between climatic and edaphic factors and primary productivity across land use types (woodlands, grasslands, and croplands). Explained variation is expressed as R^2^, and significance is indicated by asterisks (***; p-value < 0.001, **; p-value < 0.010, *; p-value < 0.050). Non-significant (p-value > 0.05) regressions are denoted with “n.s.”.

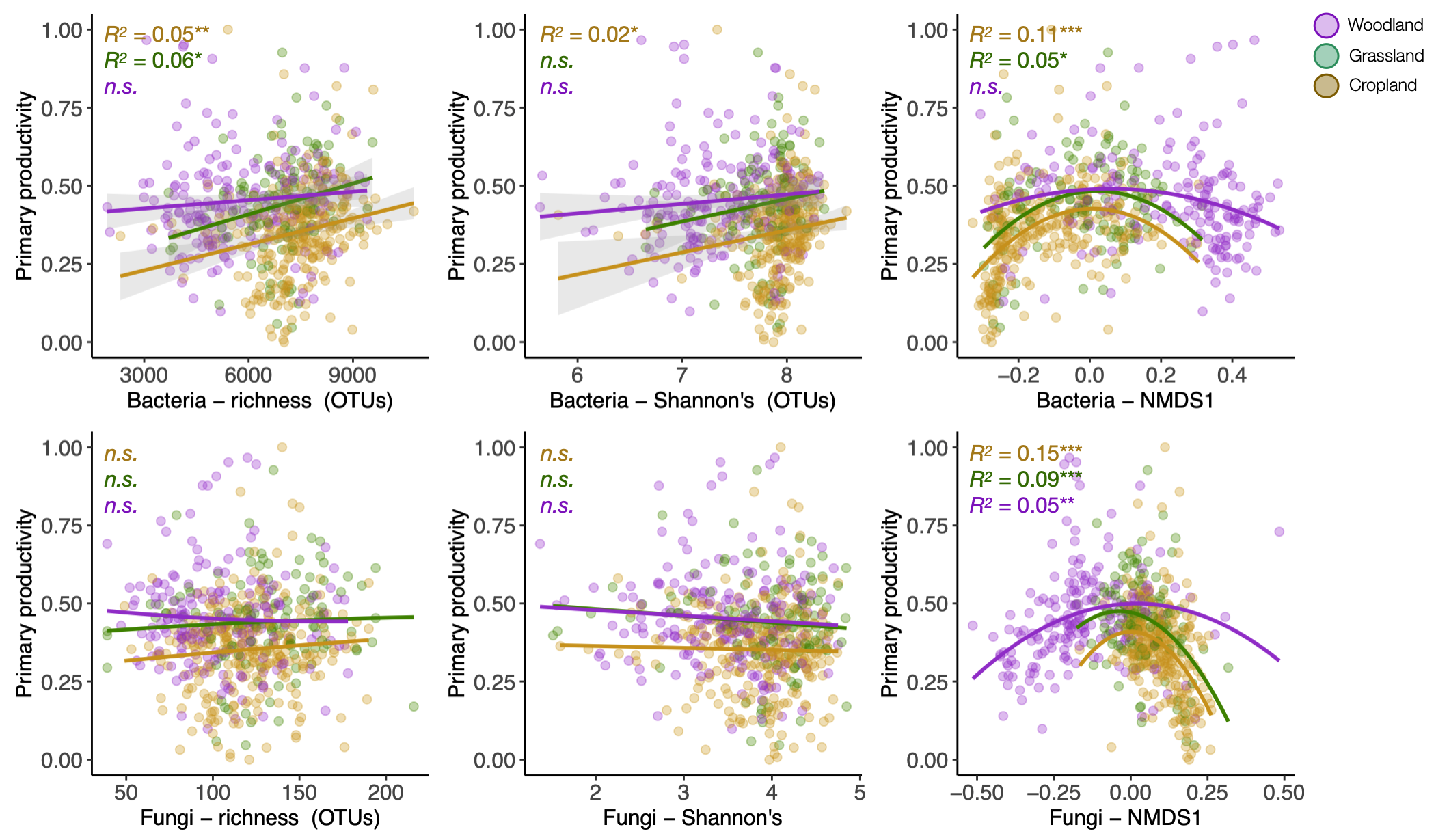

**Figure S2.** **Relationships between microbial community metrics and primary productivity**. Linear regressions (polynomial for community composition) were fitted to explore the relationship microbial community metrics including total OTU richness, Shannon’s diversity index, and community composition (NMDS first dimension) with primary productivity across land use types (woodlands, grasslands, and croplands). Explained variation is expressed as R^2^ (0-1), and significance is indicated by asterisks (***; p-value < 0.001, **; p-value < 0.010, *; p-value < 0.050). Non-significant (p-value > 0.050) regressions are denoted with “n.s.”.

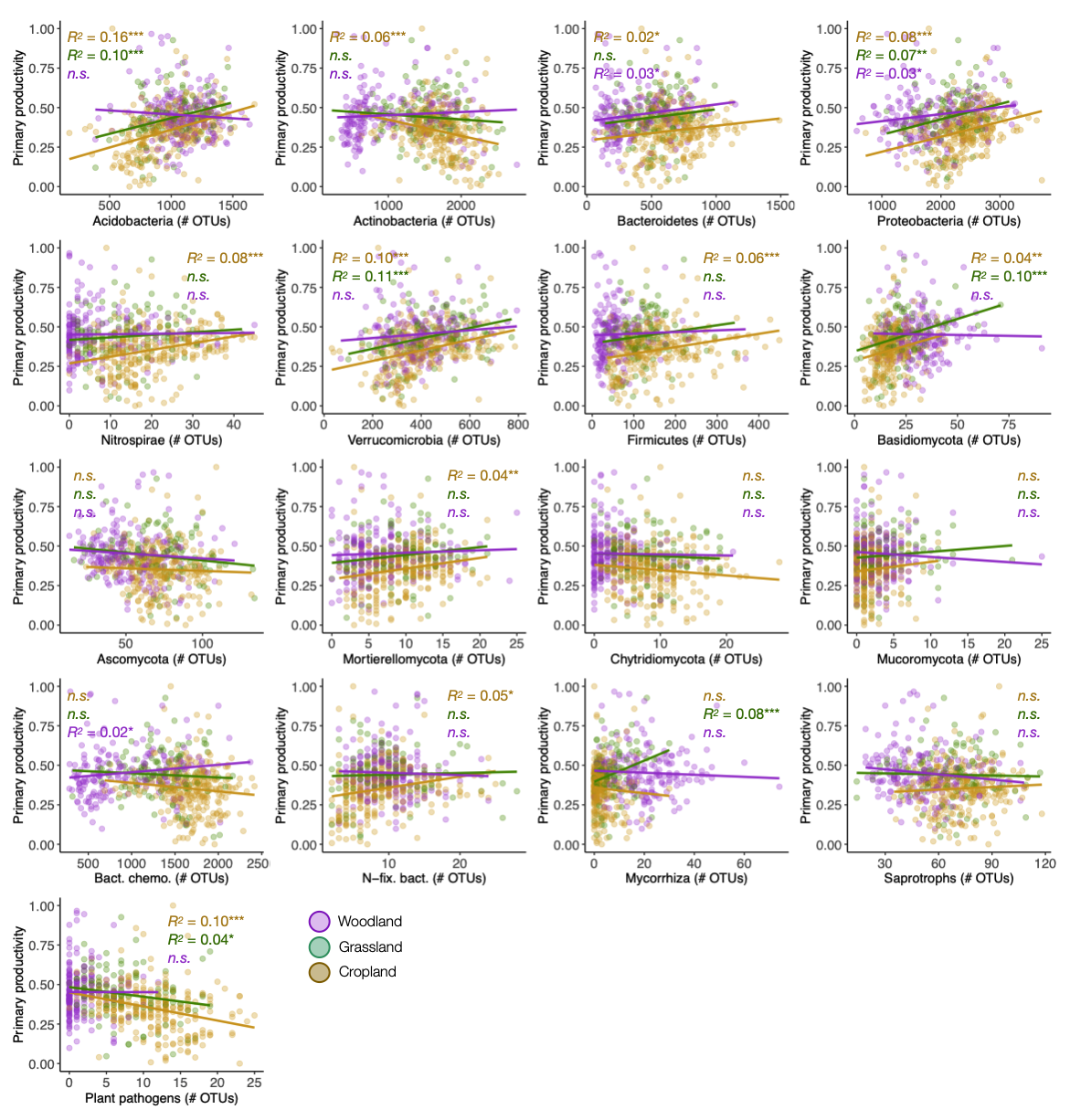

**Figure S3.** **Relationships between soil biodiversity and primary productivity.** Linear regressions were fitted to explore the relationship between the main bacterial and fungal phyla and primary productivity across land use types (woodlands, grasslands, and croplands). Linear regressions were also fitted to the relationship between primary productivity and the main functional groups (Bacterial chemoheterotrophs, N-fixing bacteria, mycorrhizal fungi, saprophytic fungi, and fungal plant pathogens). Explained variation is expressed as R^2^, and significance is indicated by asterisks (***; p-value < 0.001, **; p-value < 0.010, *; p-value < 0.050). Non-significant (p-value > 0.05) regressions are denoted with “n.s.”.

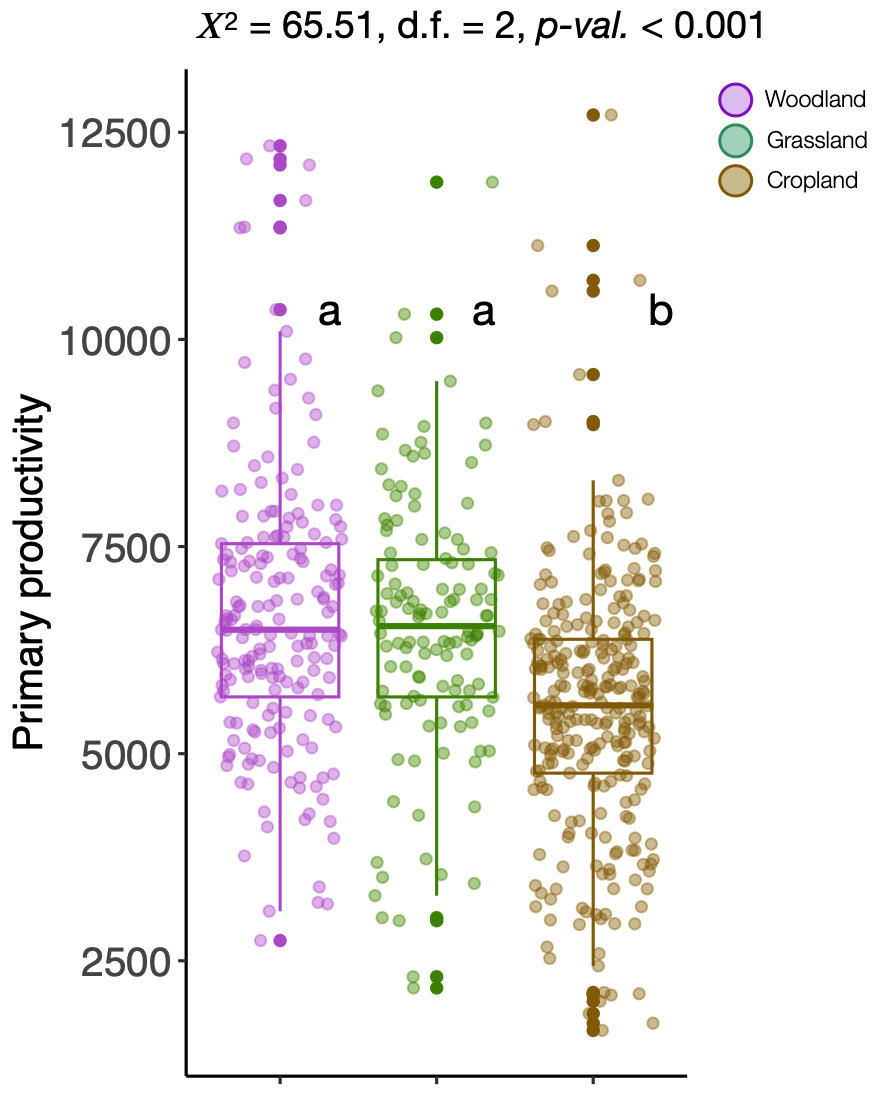

**Figure S4.** **Primary productivity across land use types.** Primary productivity data as obtained from Terra and Aqua satellites across land use types (woodlands; n = 186, grasslands; n = 126, croplands; n = 277). Grouping information (letters above boxplots) is provided pairwise comparisons following Kruskal-Wallis rank test (different letters indicate significant difference at *p* < 0.05). Chi-squared (𝜒^2^) provides an estimation of the overall difference among land use types (***; p-value < 0.001, **; p-value < 0.010, *; p-value < 0.050).

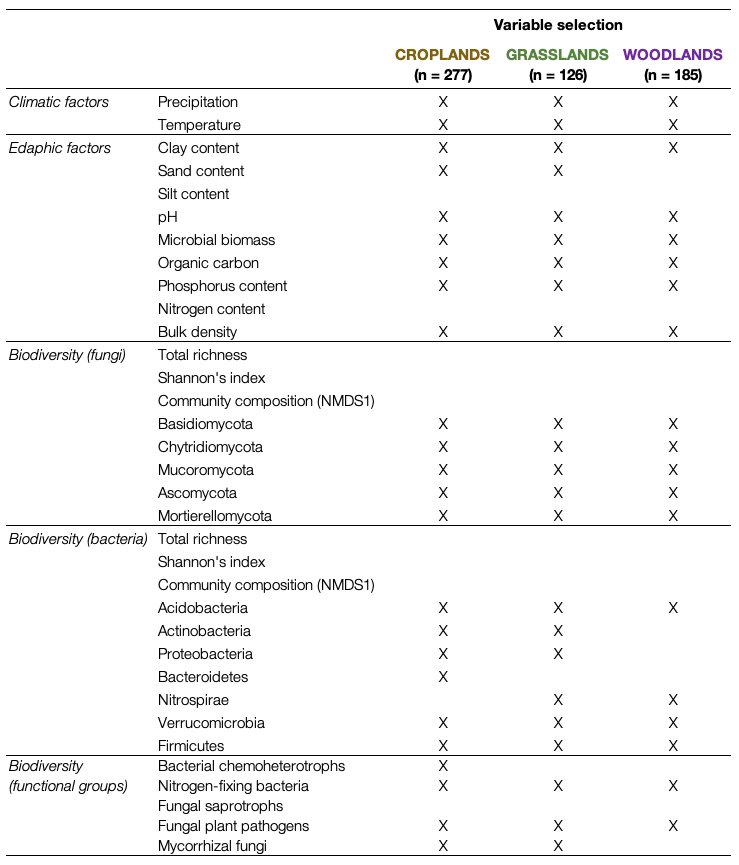

**Table S1.** Variable selection for the cropland, grassland, and woodland dataset. A total of 34 explanatory variables were tested for collinearity after applying a threshold of ±0.70 (Spearman’s rank coefficient). Variables not selected with “X” were excluded before further analyses because high multicollinearity was detected.

| **Variable** | **Reason** | **Reference** |
| --- | --- | --- |
| Nitrogen-fixing bacteria | N-fixing bacteria play a crucial role in converting atmospheric nitrogen into forms available to plants, thereby enhancing soil fertility and reducing the need for nitrogenous fertilizers | (Vitousek et al., 2013) |
| Mycorrhizal fungi | Mycorrhizal fungi form symbiotic relationships with most plants, aiding in nutrient uptake, particularly phosphorus, enhancing plant health, and improving soil structure. | (Gupta, 2020; van der Heijden et al., 2015) |
| Soil organic carbon | Soil organic carbon promotes water retention, supports microbial biodiversity, and acts as a sink for atmospheric CO_2_. | (Weil & Magdoff, 2004) |
| Phosphorus content | Phosphorus is vital for plant energy transfer and contributes to microbial functions including organic matter decomposition | (Duncan et al., 2019) |
| Soil density | Soil density is a measure of soil compaction, and influences root penetration, water infiltration, and aeration. Lower density indicates good soil structure with sufficient pore spaces. | (Shah et al., 2017) |
| Microbial biomass | Microbial biomass represents the living part of soil organic matter and is a key indicator of microbial activity | (Zhou & Ding, 2007) |
| Pathogen richness | The richness of pathogenic taxa in a soil provides a measure of disease risk and potential loss in productivity | (Liu et al., 2022) |

**Table S2.** Variables included in the soil health index.

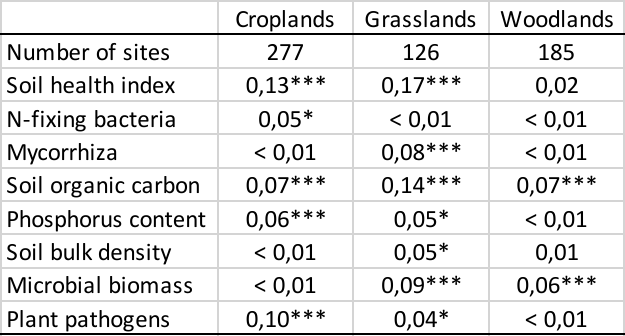

**Table S3.** Relationship between individual variables and the soil health index with primary productivity. This table presents R-squared values (0-1) reflecting the strength of association between each individual variable included in the index and primary productivity. The table also shows the correlation between the aggregated variables (i.e., soil health index) and primary productivity, allowing for a comparison between the relative performance of the soil health index against the individual variables. Significance is indicated by asterisks (***; p-value < 0.001, **; p-value < 0.010, *; p-value < 0.050).

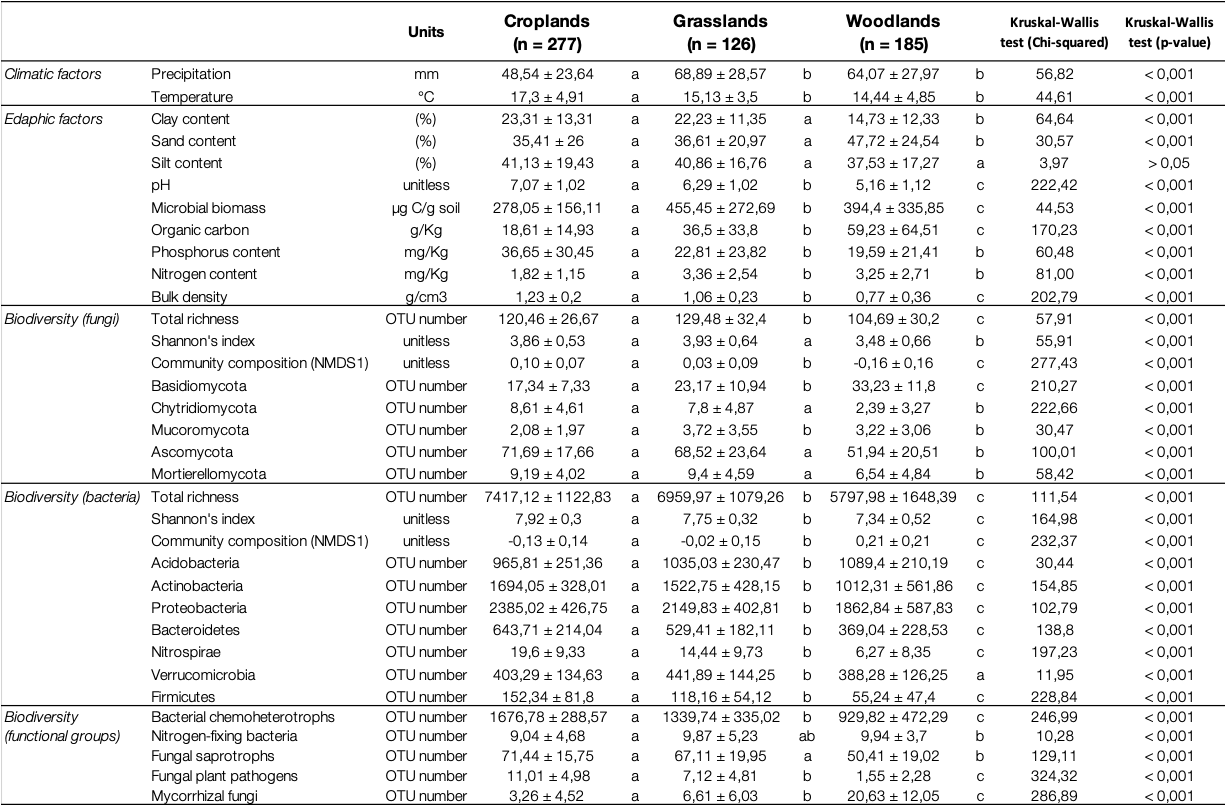

**Table S4** Mean values of spatial, climatic, and edaphic factors (± standard deviation) in soils from cropland (n = 277), grassland (n = 126), and woodland (n = 185) sites used in this study. Mean values of biodiversity measures are also shown for bacteria, fungi, and predicted functional groups. Kruskal-Wallis test was used to assess the overall effect of land use type on explanatory variables (chi-squared and p-value are indicated for each test). Grouping information following pairwise Wilcoxon test is indicated above each boxplot, different letters indicate significant differences (p < 0.05).
